# RNA interference mediates RNA toxicity with parent-of-origin effects in *C. elegans* expressing CTG repeats

**DOI:** 10.1101/2021.05.19.444826

**Authors:** Maya Braun, Shachar Shoshani, Joana Teixeira, Anna Mellul Shtern, Maya Miller, Zvi Granot, Sylvia E J Fischer, Susana M D A Garcia, Yuval Tabach

## Abstract

Nucleotide repeat expansions are a hallmark of over 40 neurodegenerative diseases. These repeats cause RNA toxicity and trigger multisystemic symptoms that worsen with age. RNA toxicity can trigger, through an unclear mechanism, severe disease manifestation in infants that inherited repeats from their mothers. Here we show in *Caenorhabditis elegans* how RNA interference machinery causes intergenerational toxicity through inheritance of siRNAs derived from CUG repeats. The maternal repeat-derived small RNAs cause transcriptomic changes in the offspring, reduce motility and shorten lifespan. However, the toxicity phenotypes in the offspring can be rescued by perturbing the RNAi machinery in affected mothers. This points to a novel mechanism linking maternal bias and the RNAi machinery and suggests that toxic RNA is transmitted to offspring and causes disease phenotypes through intergenerational epigenetic inheritance.

## Introduction

Over 40 genetic diseases are caused by an expansion of a short nucleotide repeat sequence in the genome^1^. RNAs transcribed from the expanded regions were shown to disrupt cellular function through a gain-of-function or loss-of-function type of mechanisms, causing RNA toxicity. Included in the gain-of-function mechanisms are sequestration of RNA-binding proteins, microRNA and siRNA dysfunction, and RAN translation^2–6^.

Several repeat-based disorders exhibit a correlation between disease severity and gender of the transmitting parent, also known as the parent-of-origin effect^7–9^. Maternal bias is observed in myotonic dystrophy type 1 (DM1), the most common form of adult-onset muscular dystrophy, an autosomal dominant neuromuscular disease^10,11^. DM1 is caused by an unstable CTG repeat expansion in the 3’ untranslated region (3’UTR) of a serine-threonine protein kinase gene, DMPK. In healthy individuals the DMPK gene contains 5-37 CTG repeats, whereas DM1 patients bear a range of 50 to several thousand repeat expansions^12^. Patients present a variety of symptoms but are mainly characterized by muscle dysfunction, progressive muscle wasting, atrophy, and myotonia. DM1 is classified into five subtypes based on repeat length and age of onset, marking congenital myotonic dystrophy as exceptional in parameters of severity and pathophysiology of the disease. In contrast to the other subtypes that are characterized by late onset and progressive diseases, in congenital DM1 the symptoms already appear during pregnancy, with infants displaying upon birth severe generalized weakness and respiratory insufficiency. The newborns suffer from life-threatening complications and display up to 40% mortality^8^. Contrary to expected deterioration over time, symptoms improve during childhood until adult-onset DM1 symptoms develop during adolescence^13^. Congenital myotonic dystrophy is almost exclusively of maternal transmission^14,15^. Comparison of parental inheritance shows generally worse outcomes in DM1 children born to affected mothers^16^. Moreover, non-DM1 carrier children born to DM1 mothers show a worse medical prognosis following pre-implantation genetic diagnosis^17,18^. Together, these data demonstrate a clear gender bias. While DM1 is the best characterized disease, additional expansion repeat disorders presenting an early-onset form with distinct clinical features include Huntington’s Disease-like 2 (HDL2) and spinocerebellar ataxia type 8 (SCA8)^7,19,20^. Both are caused by CUG repeats in the 3’UTR, suggesting a global mechanism driven by the toxic CUG RNAs.

The mechanism underlying this parent-of-origin toxic effect remains unclear. Upstream CpG methylation was linked with maternal bias for transmission^21^, however it does not account for the different disease phenotypes observed. Possible factors such as extended maternal repeat length, mitochondrial DNA mutations or maternal intrauterine factors have not been identified^22–26^ thus implying that the contribution of additional factors should be considered^14,27^.

RNA interference (RNAi) is a conserved gene-silencing mechanism that plays important roles in regulation of viruses, transposons and genes^28^. Genetic and biochemical analyses revealed numerous proteins (such as Argonaute and Dicer) as essential cofactors that process and present small RNAs to their targets^29–35^. The enzyme Dicer processes trigger double-stranded RNA (dsRNA) into short-interfering RNA (siRNA) of ∼21 nucleotides. These form a protein-RNA complex that degrades and inhibits translation of the target messenger RNA (mRNA) bearing a complementary sequence^36^. Systematic analysis of the RNAi pathway revealed conservation and co-evolution of the associated proteins across eukaryotes^31^. Many of the early discoveries in the field^32,34,35,37–43^ were established in *Caenorhabditis elegans* nematodes due to the availability of unique genetic tools that allowed mapping and characterization of the RNAi pathway.

The RNAi machinery is also associated with various pathological states^44–46^. Specifically, it was found to contribute to the aberrant activation of pathogenic RNA toxicity mechanisms in nematodes^47^, drosophila^48^, and mammals^49–51^. Expanded DNA repeats, when transcribed, tend to form stable hairpins that accumulate as RNA foci in the nucleus and cytoplasm^4,52–54^. Dicer can target these double-stranded RNAs with an imperfect pairing and cleave them to siRNAs^50,51,55,56^. These repeat-derived siRNAs engage the downstream RNAi silencing pathway and cause global changes in the expression of endogenous genes bearing complementary sequences^47^.

The RNAi machinery also mediates heritable epigenetic modulations, a phenomenon termed RNAi inheritance^57–68^. siRNAs are inherited from parent to progeny through oocytes and potently prompt transgenerational gene silencing^64^. Recently, a maternal parent-of-origin effect was observed in the transmission of small RNA-based endoderm development phenotypes and RNAi machinery defects^57,69,70^. In mammals, the presence of small RNAs from maternal origin was shown in human and mouse cord blood^71–75^.

Here, we use *C. elegans* to uncover the mechanisms that underlie the parent-of-origin RNA toxicity effect. We hypothesize a key role for the RNAi machinery in these pathogenic processes. Using nematodes has the advantage of providing an isogenic system to address these mechanisms in animals, which enables efficient perturbation of investigated genes and consequent examination of phenotypic, developmental, and cross-generational changes. We recapitulate a maternal bias in a *C. elegans* model expressing expanded CUG repeats and show that disrupting the RNAi pathway in affected mothers rescues toxicity in offspring. Overall, our work points to the central role of the RNAi pathway as the underlying mechanism of the parent-of-origin effect present in RNA repeat-based pathogenesis.

## Results

### *In vivo* isogenic model for studying phenotypes of long RNA repeats

We adapted a *C. elegans* experimental system^47^ to identify the mechanisms underlying parent-of-origin toxicity associated with CUG repeat expansions, specifically the potential role of RNAi machinery in these toxic processes. We used *C. elegans* strains expressing GFP containing either 0 or 123CTG repeats (0CUG and 123CUG, respectively) in the 3’UTR, under the regulation of the *myo-3* muscle-specific promotor (Fig. 1A)^47,52^. Expression of 123CTG repeats was shown to be sufficient to induce RNA toxicity in mammalian myocytes and nematodes, whereas longer repeat lengths resulted in embryonic lethality^47,52,76^. To exacerbate the toxicity induced by the repeat-bearing RNAs, transgenic strains were generated expressing several copies of the repeats as an extrachromosomal array integrated in the genome (see Discussion).

**Figure 1:**
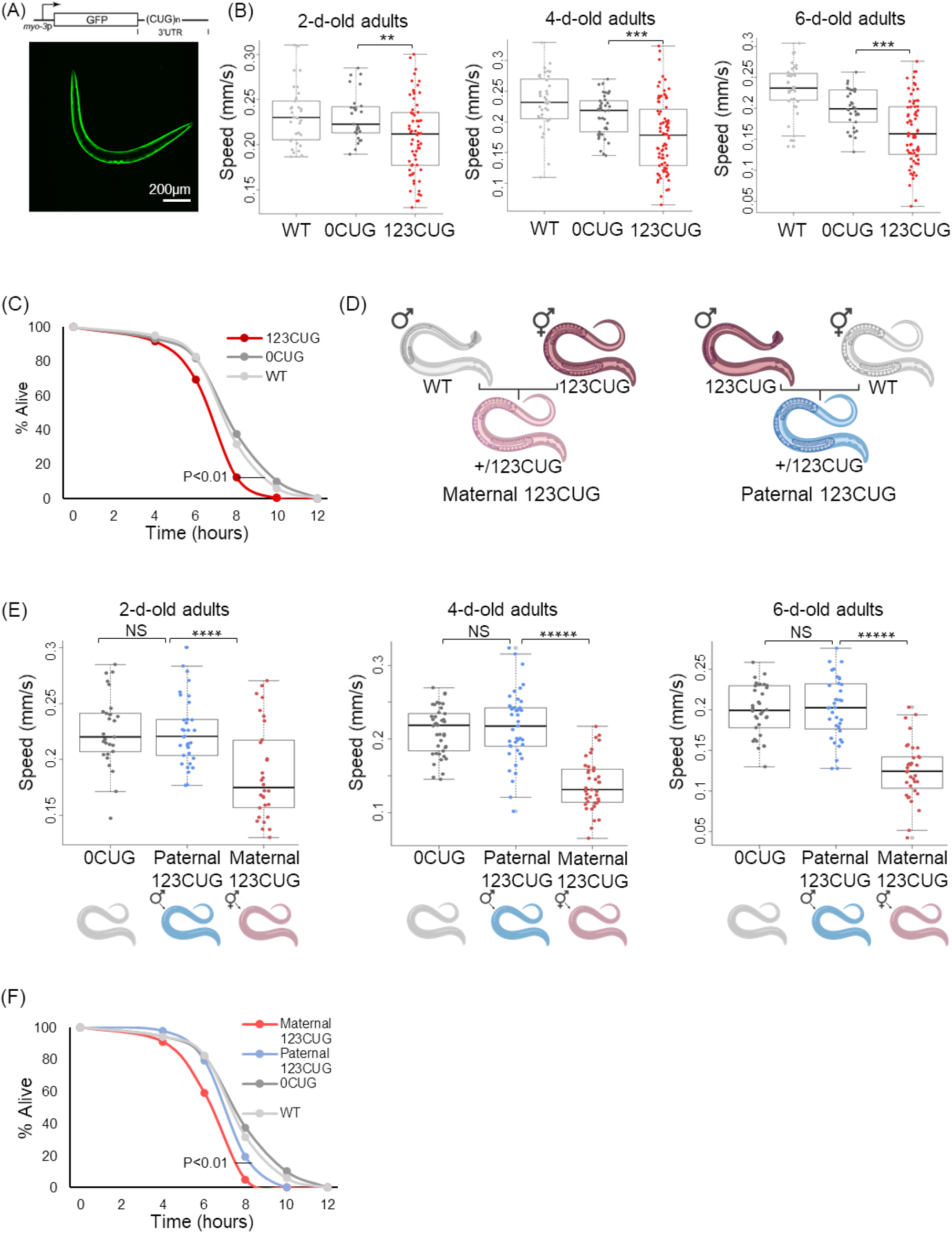
Maternal inheritance of repeats aggravates disease phenotype. (A) *C. elegans* model for repeat expansions. 123 CUG repeats are expressed in the 3’UTR of GFP under the *myo-3* promoter. Representative image of a 123CUG nematode. (B) Motility assay (moving speed) of WT, 0CUG, and 123CUG nematodes (n=60) for two-, four-, and six-day-old adults. (C) Survival curve of WT, 0CUG, and 123CUG day one adult nematodes following heat shock (35°C, n=80). (D) A general scheme of the parent-of-origin experimental system. (E) Motility assay for assessment of parent-of-origin effects. Moving speed of 0CUG, paternally inherited 123CUG, and maternally inherited 123CUG nematodes (n=45). (F) Survival curve of Maternal 123CUG and Paternal 123CUG animals following heat shock (35°C, n=80). In the motility assays (B and E), the average of three biological replicates is represented. Data are represented as a mean ± SD and significance was calculated using a one-tailed Student’s t test. In the heat shock assays (C and F), the results shown are from a representative experiment of three biological replicates. Statistical analyses were performed using log-rank (Mantel-cox) and Gehan-Breslow-Wilcoxon tests. * p<0.05; ** p<0.01; *** p<0.00l; **** p<0.000l: NS - not significant.

To reconfirm the robustness of the model, we quantified the baseline toxicity of the expanded repeats^47^. We conducted a motility assay and measured the speed of movement of the nematodes using video recording under light microscopy at three different ages (2^nd^, 4^th^, and 6^th^ day of adulthood). The average moving speed of 0CUG nematodes was 0.231 mm/s, 0.21 mm/s, and 0.199 mm/s at 2, 4, and 6-day-old adults. 123CUG animals moved at an average speed of 0.209 mm/s, 0.178 mm/s, and 0.164 mm/s at 2, 4, and 6-day-old adults, exhibiting impaired motility that deteriorated with age (Fig. 1B)^47^. Next, we examined how the repeats affect susceptibility to stress. We exposed wild type, 0CUG, and 123CUG nematodes to heat shock (35°C) and scored for viability every two hours. 123CUG animals showed a higher susceptibility to heat stress (p<0.01, Fig. 1C).

### Maternal inheritance of CUG repeats enhances motility impairment and susceptibility to stress

To investigate the parent-of-origin effect, we assessed toxicity in offspring with either maternal or paternal inheritance of repeats. We generated 123CUG males and hermaphrodites. We crossed 123CUG males with wild type hermaphrodites and 123CUG hermaphrodites with wild type males to assess paternal (Paternal 123CUG) and maternal (Maternal 123CUG) transmission of repeats, respectively (Fig. 1D). The 123CUG animals did not show changes in repeat size in successive generations (Table 1S). Thus, the Maternal and Paternal 123CUG offspring share the same genetic background, expressing transcripts with the same repeat length (heterozygous 123CUG). Next, to determine whether differences in toxicity were present, we performed motility and heat stress assays and compared the two groups.

Maternal inheritance of repeats resulted in severely impaired motility. Animals showed an average moving speed of 0.188 mm/s in two-day-old Maternal 123CUG adults as compared with 0.225 mm/s in the 0CUG group (p<0.0001). Importantly, the motility of animals with paternal inheritance of toxic repeats was indistinguishable from the 0CUG controls, with a speed of 0.22 mm/s (Fig. 1E). We then tested these animals for changes in motility with aging and observed an age-dependent gradual loss of motility.

The Maternal 123CUG animals displayed speeds ranging from 0.188 mm/s to 0.125 mm/s from the 2^nd^ to the 6^th^ day of adulthood, respectively. Thus we chose two-day-old adults for the following assays as they have a strong phenotype and are less susceptible to confounding environmental factors. Next, to check whether the maternal bias is reproduced in additional phenotypes, we exposed the animals to heat shock (Fig. 1F). Maternal 123CUG nematodes were significantly more susceptible to heat stress, with survival rates of 55% after six hours as compared with 80% in the control groups. After eight hours, only 5% of Maternal 123CUG nematodes survived as compared with 18% of Paternal 123CUG and over 30% in the control groups. Based on the hypothesis of asymmetric inheritance of toxicity, we tested whether an effect on normal nematode growth was present. We assessed nematode morphology and found that the Maternal 123CUG animals were significantly shorter at the 1^st^ larval stage, with an average length of 173.2 μm as compared with 187.5 μm in the 0CUG animals (p=1.83*10^−6^). Notably, the length of the Paternal 123CUG nematodes was similar in size as the 0CUG controls. Aging led to a reduction in the gap – the length was 1327 μm in two-day-old adults Maternal 123CUG as compared with 1368 μm in Paternal 123CUG and 1382 μm in 0CUG (p=0.002, Fig. 1S). To eliminate a possible strain-specific bias, which might result from the location site of repeat integration in the genome or an uncontrolled evolutionary effect such as selection or random drift, we repeated the motility experiment with different 0CUG and 123CUG strains (GR3208 and GR3207, respectively). These experiments supported the previous results and exhibited an enhanced maternal effect (Fig. 2S).

**Figure 2:**
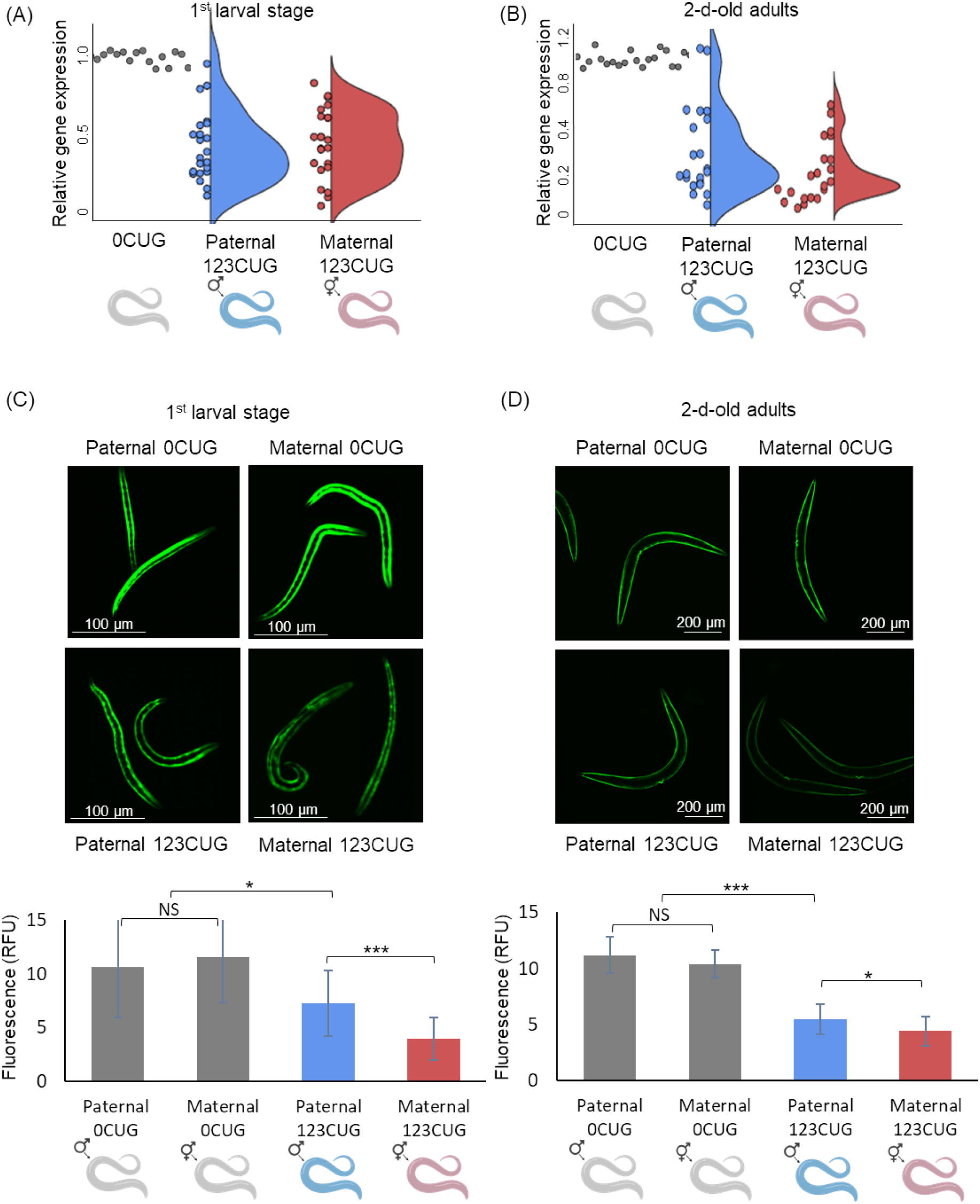
Maternal inheritance of repeats escalates downregulation in expression of genes bearing endogenous CTG- and CAG-repeats. Gene expression fold change of 24 genes bearing ≥ 7 CTG/CAG repeats in L1 (A) and 2-day-old adults (B), relative to 0CUG. The qPCR is an average of 3 biological experiments and 3 technical replicates. Fluorescence levels of Maternal 0CUG, Paternal 0CUG, Maternal 123CUG, and Paternal 123CUG nematodes in (C) 1^st^ larval stage and (D) 2-day adults. Relative fluorescence was computationally quantified (n=50) and representative fluorescent microscopy images are shown. Data are represented as a mean ± SD of three biological replicates and significance was calculated using a one-tailed Student’s t test. * p<0.05, *** p<0.00l.

### Maternal inheritance of repeats suppresses gene expression of genes bearing endogenous CTG- and CAG-repeats

Across species, transcribed CUG- and CAG-repeats trigger abnormal gene silencing induced by the RNA interference^47,50,51^. In both humans and *C. elegans* expressing expanded repeats, silencing was specifically enriched in genes bearing complementary short repeat sequences, a hallmark of silencing by the RNAi machinery^47^. We aimed to test whether the RNAi pathways play a role through a similar mechanism in the parent-of-origin toxicity effect.

First, we determined that the CUG repeats were targeted by the RNAi machinery and processed to siRNAs. We measured the levels of siRNAs comprised of seven CUG repeats in Paternal 123CUG and Maternal 123CUG two-day-old adult nematodes and compared them with wild type nematodes (see Methods). Both Maternal and Paternal 123CUG animals showed an enrichment of repeat-derived siRNAs (Fig. 3S). Next, we checked if there is a difference in siRNA-dependent silencing between the two groups, suggestive of the involvement of the pathway in parent-of-origin toxicity. We have previously shown through RNAseq transcriptome analysis of 123CUG animals, that a broad decrease in expression levels is observed in endogenous genes bearing CTG repeats. The expression of these genes was enriched in muscles, neurons, hypodermis, and intestine, correlative to the presented disease phenotypes. Here, we measured the expression levels of 24 potential targetable genes that contained seven or more endogenous CTG or CAG repeats and were previously validated as markers of CTG-targeted silencing^47^. Gene expression was compared between offspring with Maternal and Paternal 123CUG. We collected RNA from the 1^st^ larval stage and two-day-old adult nematodes expressing Maternal 123CUG, Paternal 123CUG, or 0CUG. Using qPCR, we analyzed the expression of the 24 genes. Two-day-old adult Maternal 123CUG animals showed a significant 4-fold down-regulation in expression, with Paternal 123CUG displaying a 2.6-fold reduction, both relative to the 0CUG controls (Fig. 2A-B). These strains were analyzed at a younger developmental stage, i.e. the 1^st^ larval stage, and exhibited a slightly more severe gene silencing phenotype. The Maternal 123CUG group exhibited a 2-fold silencing relative to 0CUG, whereas the Paternal 123CUG group showed a 1.6 down-regulation (Fig. 2A-B). Overall, these genes present stronger repression in progeny with Maternal 123CUG as compared with Paternal 123CUG.

**Figure 3:**
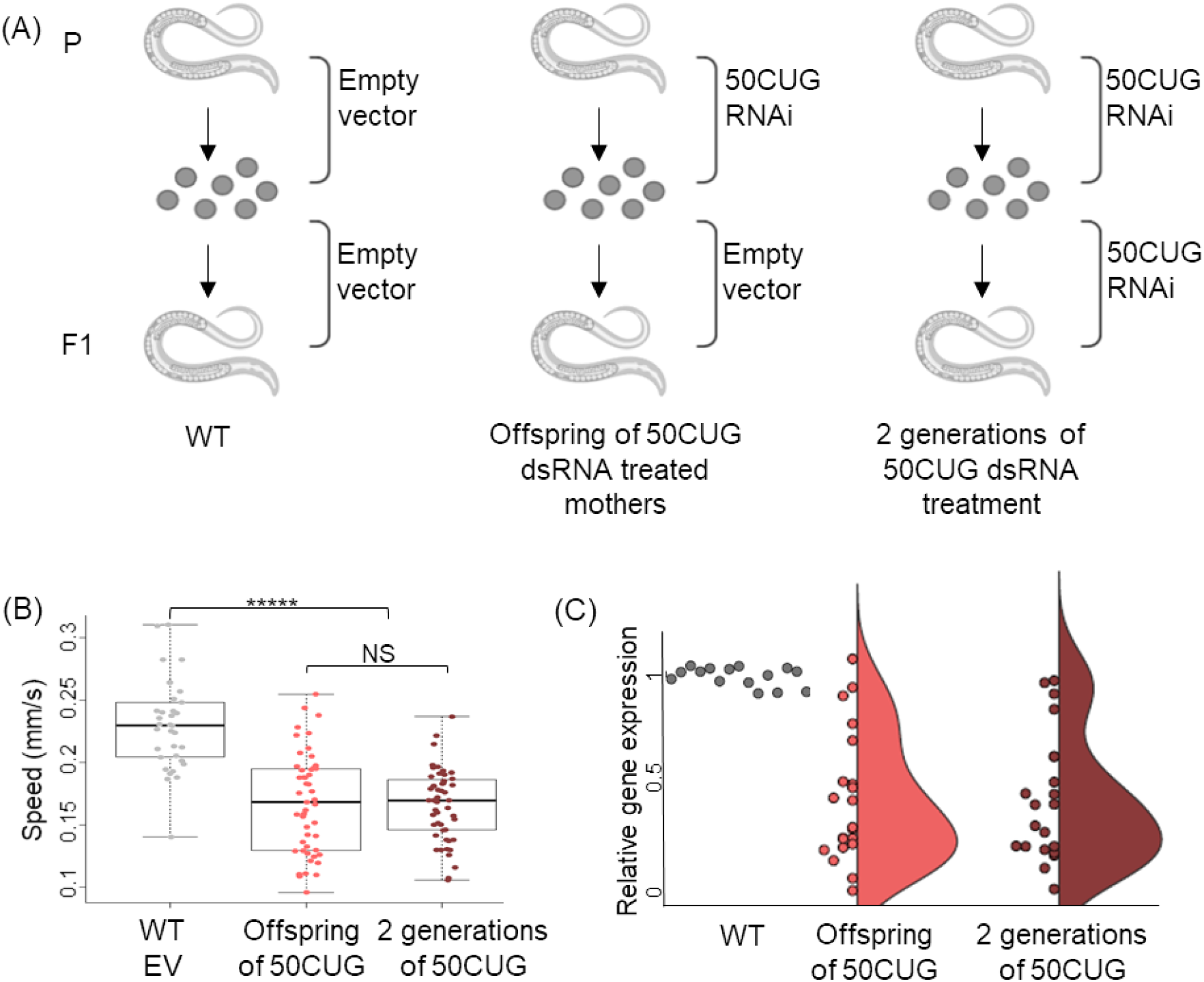
Feeding nematodes with 50CUG dsRNA recapitulates toxicity phenotypes. (A) Experimental scheme: Wild-type nematodes were fed dsRNA bearing 50CUG repeats for one or two generations. (B) Motility assay of two-day-old adult WT, offspring of nematodes treated with 50CUG dsRNA, and nematodes fed 50CUG dsRNA for two generations (n=45). Data are represented as a mean ± SD of three biological replicates and significance was calculated using a one-tailed Student’s t test. Box plots: center line, median; box limits, upper and lower quartiles. ***** p<0.00001, NS - not significant. (C) Gene expression fold change of 24 genes bearing ≥7 CTG/CAG repeats in two-day-old adults as determined by qPCR for 50CUG treatment groups relative to wild type. Three biological and three technical replicates were analyzed for this experiment.

In addition to transcriptomic changes, 123CUG animals show a noticeable decrease in GFP protein levels with age^47,52^ as compared with 0CUG, which may point to transgene silencing^77,78^. To establish whether the parental origin of repeats causes variability in GFP levels, we measured and compared the fluorescence levels of Maternal 123CUG, Paternal 123CUG, Maternal 0CUG, and Paternal 0CUG nematodes at two different life stages (1^st^ larval stage and two-day-old adults). The Maternal 123CUG animals exhibited a 2-fold stronger decay in fluorescence as compared to the Paternal 123CUG animals at the 1^st^ larval stage (Fig. 2C). This trend persisted, although attenuated, for two-day-old adults (Fig. 2D). There was no difference between the Maternal 0CUG and Paternal 0CUG strains. Together these data indicate that the RNAi machinery processes the expanded repeats to siRNAs and mediate aberrant silencing of genes bearing complementary sequences.

### Directly activating the RNAi pathway by feeding nematodes 50CUG dsRNA recapitulates toxicity phenotypes

The integrated repeats in the 123CUG nematodes may have various uncharacterized effects as a consequence of how transgenes are generated in *C. elegans*. Although the previous experiments were repeated in two independent strains, we wished to take a complementary approach in dissecting the explicit effect of the repeats. Consequently, we directly triggered the RNAi machinery using exogenous dsRNA bearing expanded repeats. This approach enables the accurate determination of the phenotypic effects of the repeats, as they serve as direct substrates to the RNAi machinery and therefore exclude other confounders mediating RNA toxicity mechanisms. To this end, we fed wild type animals with bacteria expressing RNAs bearing 50CUG repeats (50CUG RNAi). We assessed three groups: nematodes grown on an empty vector (EV), offspring of hermaphrodites fed 50CUG dsRNA, and nematodes fed 50CUG dsRNA for two generations (Fig. 3A).

Two generations of 50CUG dsRNA-treated animals (P0 and F1 were treated) and offspring of treated animals (F1 were not treated) all exhibited impaired motility with an average moving speed of 0.166 mm/s and 0.168 mm/s, respectively, as compared with 0.229 mm/s in the wild type group grown on EV (p=1.18*10^−10^, Fig. 3B). A similar effect of suppression was observed in the expression of the 24 CTG/CAG-containing genes (as defined above). These target genes were down-regulated on average by 3.7 fold in the offspring of 50CUG group, and 3.5 fold in the group that received 50CUG treatment for two generations, relative to the wild type group (Fig. 3C). The offspring of 50CUG dsRNA treated mothers were similarly impaired in animals that were directly treated with dsRNA containing expanded repeats.

### Altering the RNAi machinery rescues toxicity in offspring with maternally inherited repeats

As both the 50CUG dsRNA and the integrated 123CUG repeats had a toxic effect on P0 and F1 generations, we tested whether motility defects, sensitivity to stress, and downregulation of the 24 target genes could be rescued. Specifically, we considered that if the RNAi machinery plays a central role in the toxicity effect observed in maternal versus paternal inheritance, then disruption of the RNAi pathway has the potential to revert the phenotypes in the F1 progeny and affect the differences in toxicity between maternal and paternal inheritance.

As knocking out key players in the RNAi machinery can cause severe phenotypes in nematodes and other animals^78,79^, we used RNAi to regulate components in the RNAi machinery. This well-validated approach partly silences proteins in the pathway, thus affecting the steady state of the RNAi machinery, but generally with limited effects on phenotype^31,37^. Complete knockout of *dcr-1* and *rde-4* RNAi resulted in severely impaired animals (Fig. 4S) whereas knockdown using RNAi^31,37^ did not produce an obvious detrimental effect (Fig. 4, 5S). Consequently, Maternal 123CUG (offspring of 123CUG mothers) were treated with RNAi against key members in siRNA production: *dcr-1, rde-4*, and *rde-1* (Fig. 4A). The treated Maternal 123CUG nematodes exhibited rescued motility, with speeds corresponding to an average of 0.22 mm/s following treatment of *dcr-1, rde-1*, and *rde-4* RNAi as compared to the empty vector (Fig. 4B, 5S). Treated Maternal 123CUG animals also exhibited improved heat shock response, with approximately 90% survival rates after six hours in 35°C as compared with 60% of Maternal 123CUG and 70% of Paternal 123CUG animals (Fig. 4C). Next, we used qPCR to assess the effect of RNAi treatment on the expression of the targeted CTG/CAG-bearing genes. We observed a 4-fold reduction in expression of targeted genes in the Maternal 123CUG group as compared to a reduction of approximately 1.3-fold following treatment with *dcr-1, rde-1*, and *rde-4* RNAi in Maternal 123CUG animals. In sum, RNAi treatment suppressed the phenotype at the molecular level and partly rescued down-regulation of the CTG/CAG-bearing genes (Fig. 4D). In comparison with the Maternal 123CUG group, an average of a 120% increase in GFP intensity was observed in the *dcr-1, rde-1*, and *rde-4* RNAi treatment groups (Fig. 4E, 6S). Notably, fluorescence levels after treatment were similar to the Paternal 123CUG group levels. Together these data support, for the first time, the role of siRNAs as the underlying mechanism in the parent-of-origin effect. Furthermore, they demonstrate that regulation of this pathway can rescue both molecular and organism level phenotypes of maternally inherited toxicity, as observed in Maternal 123CUG animals.

**Figure 4:**
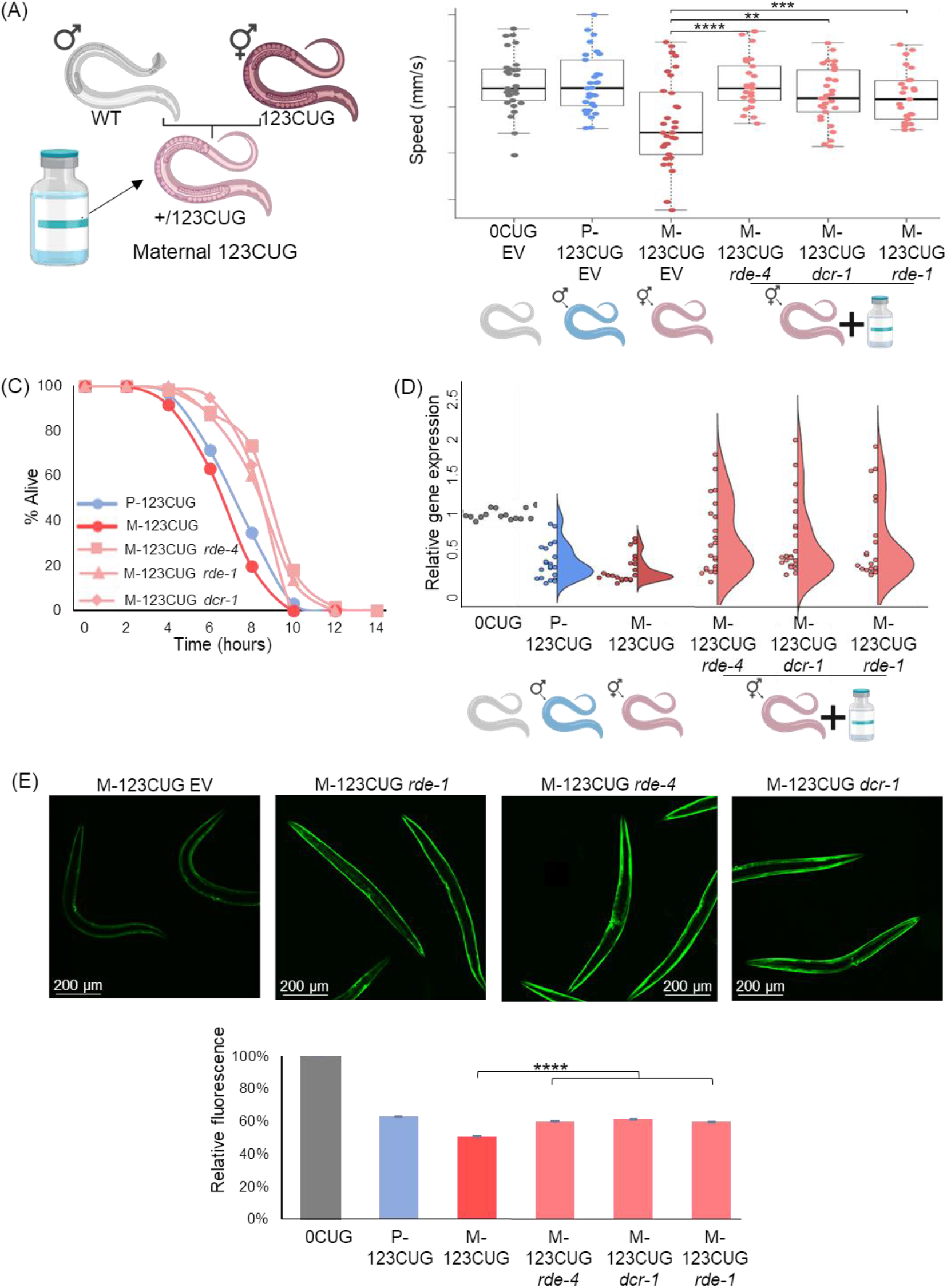
RNAi machinery is essential for maternal inheritance of repeat phenotype. (A) Scheme of the intervention: treatment of Maternal 123CUG offspring using RNAi against *dcr-1, rde-4*, and *rde-1*. (B) Motility assay of two-day-old adult Maternal 123CUG nematodes following RNAi treatment against *rde-4, dcr-1*, and *rde-1* (n=45). Data are represented as a mean ± SD of three biological replicates and significance was calculated using a one-tailed Student’s t test. (C) Survival curve under heat shock (35°C) of one-day-old adult Maternal 123CUG following RNAi treatment. The results shown are from a representative experiment of three biological replicates. Statistical analyses were performed using log-rank (Mantel-cox) and Gehan-Breslow-Wilcoxon tests (n=80). (D) Gene expression fold change of 24 genes bearing ≥7 CTG/CAG repeats; determined by qPCR for two-day-old adult Maternal 123CUG RNAi treatment groups relative to 0CUG. Three biological and three technical replicates were analyzed for this experiment. (E) Fluorescence levels of two-day-old adult Maternal 123CUG nematodes treated with RNAi. Relative fluorescence was computationally quantified (n=50) and representative fluorescent microscopy images are shown. Data are represented as a mean ± SD of three biological replicates and significance was calculated using a one-tailed Student’s t test. * p<0.05, ** p<0.01, *** p<0.001, **** p<0.0001, NS - not significant. M-123CUG, Maternal 123CUG; P-123CUG, Paternal 123CUG; M-123CUG *rde-4*, Maternal 123CUG following RNAi treatment against *rde-4*

### Reducing activity of essential RNAi machinery components in 123CUG hermaphrodites rescues toxicity in offspring

Enhancement of silencing is dependent on repeats being generated in the mothers either by feeding or expression of a transgene (Figs. 1-3). Consequently, we checked whether silencing the RNAi machinery has cross-generational effects. Epigenetic inheritance of small RNA is established^58,61,70^, but a role for this inheritance in RNA toxicity or a link to expanded repeats is unknown, as well as its potential therapeutic impact in this context. Thus, we tested if perturbing the siRNA production only in the 123CUG mothers reduced the toxic phenotype in offspring expressing 123CUG. We fed 123CUG hermaphrodites RNAi against *rde-1, rde-4*, and *dcr-1* to reduce processing of toxic RNAs into siRNAs. At the 4^th^ larval stage, we removed the hermaphrodites from the RNAi plates, crossed them with wild type males on EV plates, and assessed the offspring for effects on motility, heat shock, fluorescence, and changes in targeted gene expression (Fig. 5). Timing the RNAi feeding restricted the silencing of the RNAi machinery components (either *rde-1, rde-4*, and *dcr-1)* to the 123CUG parent hermaphrodites^64^. We observed that the progeny of 123CUG mothers with suppressed RNAi machinery displayed a significant improvement in movement with an average speed of 0.22 mm/s, as compared to 0.186 mm/s in Maternal 123CUG (Fig. 5B, 7S). A significant improvement in response to heat stress was also observed (Fig. 5C). Expression levels of CTG/CAG-bearing genes were up-regulated in the offspring of the treated mothers, from a down-regulation of 4-fold in Maternal 132CUG to 1.6-fold, and 1.3-fold in offspring of *rde-1* and *rde-4* RNAi treated mothers, reaching wild type levels following treatment against *dcr-1* in hermaphrodites (Fig. 5D). This trend was also mirrored by the fluorescence assay as fluorescence levels were on average 171% higher in the offspring of *rde-1, rde-4*, and *dcr-1* treated Maternal 123CUG nematodes in comparison to Maternal 123CUG animals (Fig. 5E). Importantly, there was no effect observed on the fluorescence levels of offspring of treated Paternal 123CUG nematodes (Fig. 8S). To conclude, down-regulating siRNA production in 123CUG hermaphrodites rescued pathogenic phenotypes in their offspring (Maternal 123CUG), but analogous treatment of 123CUG males did not affect their offspring (Paternal 123CUG). Together, these results indicate a possible treatment venue for repeat expansion disorders.

**Figure 5:**
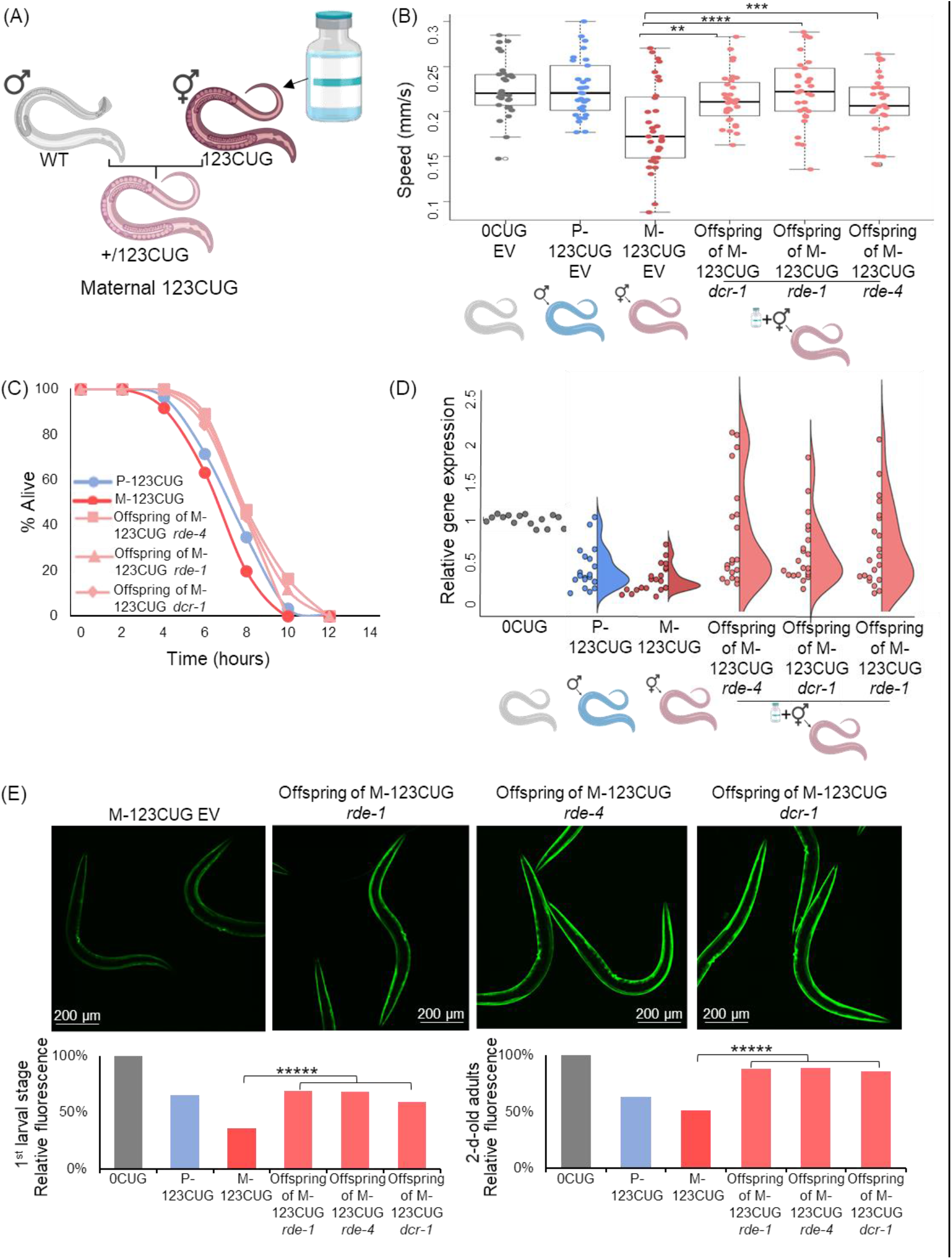
Exclusive treatment of affected 123CUG mothers rescues phenotype in offspring with maternally inherited repeats. (A) Scheme of the intervention: treatment of 123CUG hermaphrodites using RNAi against *dcr-1, rde-4*, and *rde-1*, until the 4^th^ larval stage, followed by mating with N2 males to create Maternal 123CUG animals. Offspring were observed for phenotypic effects. (B) Motility assay of two-day-old adult progeny of mothers treated against *rde-4, dcr-1*, and *rde-1* (n=45). Data are represented as a mean ± SD of three biological replicates and significance was calculated using a one-tailed Student’s t test. (C) Survival curve under heat shock (35°C) of one-day-old adult progeny of treated mothers. The results shown are from a representative experiment of three biological replicates. Statistical analysis was performed using log-rank (Mantel-cox) and Gehan-Breslow-Wilcoxon tests (n=80). (D) Gene expression fold change of 24 genes bearing ≥7 CTG/CAG repeats; determined by qPCR for offspring following RNAi treatment to mothers, relative to 0CUG. Three biological and three technical replicates were analyzed for this experiment. (E) Fluorescence levels of 2^nd^ larval stage and two-day-old adult Maternal 123CUG offspring subsequent RNAi treatment of mothers. Relative fluorescence was computationally quantified (n=50). Representative fluorescent microscopy images of two-day-old adults are shown. Data are represented as a mean ± SD of three biological replicates and significance was calculated using a one-tailed Student’s t test. ** p<0.01, *** p<0.00l, **** p<0.0001, NS - not significant. M-123CUG, Maternal 123CUG; P-123CUG, Paternal 123CUG

## Discussion

**We offer a new mechanism for the intergenerational toxicity of CUG repeats. We demonstrated how the repeated RNAs can cause maternal bias through the RNA machinery. This led to our ability to rescue most of the disease phenotypes in offspring by targeting the RNAi machinery in the mothers. Overall, we characterized in C. *elegans* a complex molecular crosstalk between DNA repeats, RNA toxicity, and the RNAi machinery in gender-specific toxicity inheritance, and its consequences to disease onset and progression**. We showed that the maternal bias associated with expanded CTG repeats is mediated by intergenerational transmission of maternal repeat-derived siRNAs that initiate early alterations in gene expression.

Generally, the parent-of-origin effect is grossly attributed to three possible mechanisms: epigenetic regulation of gene expression, mitochondrial genome mutations, and the maternal intrauterine environment^9^. While a global mechanism underlying the parent-of-origin effect in repeat expansion disorders remains unelucidated, several explanations have been proposed for the maternal bias in congenital DM1: hypermethylation upstream to the expanded repeats that alters the expression of the downstream *SIX5* gene, and longer repeat expansions leading to increased disruption of MBNL activity^21,27,80,81^. *SIX5* loss of function in model mice did not recapitulate congenital DM1 phenotypes^82^. *Mbnl1;Mbnl2;Mbnl3* triple-knockout mice have shown congenital myopathy, respiratory insufficiency, and perinatal lethality^81^. However, the correlation is limited between a larger expansion of repeats leading to increased RNA toxicity and manifestation of congenital DM1^14^. Therefore, additional mechanisms are expected to contribute to the maternal bias in CDM1 pathogenesis.

Here we established an isogenic animal system to study repeat-induced toxicity across generations. *C. elegans* are ideal to study epigenetic inheritance^58,60,83^ and the RNAi machinery^29,32,34,35,37–42^ in a well-controlled background. While our work uses *C. elegans*, the phenotypic similarity and the conservation of the RNAi machinery across eukaryotes^31^ suggests that homologous mechanisms can be found in other species. While further studies are needed to confirm the mechanism in mammals, cell lines are inappropriate for investigation of parental environment impact on progeny.

To dissect the RNA toxicity effect, we expressed 123CUG repeats endogenously, isolated from its gene context, and in addition examined the effect directly using repeat dsRNA (which we denoted as 50CUG). We found that expanded CUG repeats induced RNA toxicity with maternal biases and that this effect was dependent on the RNAi machinery (Fig. 6). We found elevated expression of small repeat RNA and coordinated suppression of genes with complementary sequences in both Maternal 123CUG (offspring of 123CUG mothers) and Paternal 123CUG (offspring of 123CUG fathers) animals, with enhanced suppression in the Maternal 123CUG group. In addition, the phenotypes associated with toxicity were asymmetrically inherited and enhanced in the Maternal 123CUG animals. Altering fundamental RNAi machinery members rescued the disease phenotypes and up-regulated the expression of affected genes. Moreover, down-regulating key members in siRNA production exclusively in the affected hermaphrodites rescued all the molecular and organismal phenotypes in the untreated offspring. These data imply that the parent-of-origin effect is mediated by maternal repeat-derived siRNAs that enhance early gene silencing in their progeny. These results point to a potential therapeutic approach for repeat-carrying mothers to ameliorate disease phenotype in progeny.

**Figure 6:**
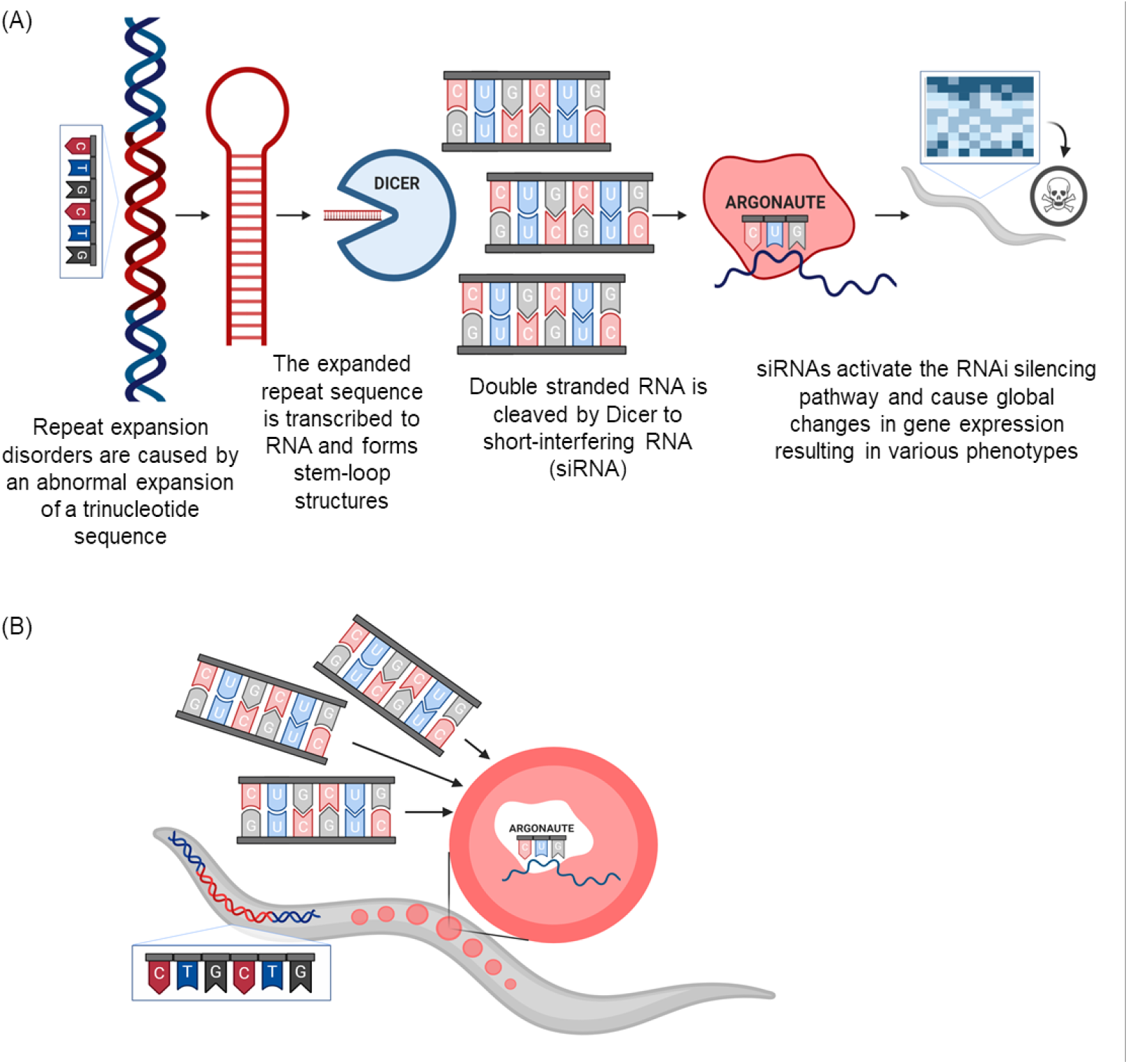
Proposed mechanism. (A) The RNAi machinery mediates RNA toxicity in expansion repeat disorders and causes maternal bias. (B) Repeat-derived siRNAs from affected mother initiate early RNAi induced toxicity in offspring

Regarding the limitations of the 123CUG system, it could be argued that the repetitive arrays are independently targeted by the RNAi machinery and processed into siRNAs. However, the recapitulation of disease phenotype achieved by directly feeding 50CUG dsRNA to nematodes, together with the rescue effect following perturbation of *dcr-1, rde-1*, and *rde-4*, strongly support the role of Dicer in mediating the toxic siRNA production. Importantly, the control data obtained from the 0CUG strain show that the maternal bias is specific to the expanded CUG repeats and is absent in the control 0CUG strain (Fig. 2B, 2S). Therefore, if transgene silencing contributes to siRNA production, it is assumed to be negligible, but nevertheless augments the desirable outcome of recapitulating the pathologic siRNAs found in patient cells.

The human genome consists of over 800 genes bearing endogenous 6CTG repeats that may serve as potential targets for siRNA silencing. Among these genes are numerous transcription factors, whose alteration may trigger extensive changes in gene expression, affecting various tissues. Future work analyzing patient transcriptome in affected tissues may shed light on unanswered questions in disease pathology. Our suggested mechanism, if recapitulated in mammals, has the potential to explain several phenotypes associated with congenital DM1 as well as other expansion repeat disorders that exhibit prominent maternal biases^7,8^. We highlight the embryonic environment as a major contributor to the aggravated phenotype associated with maternal transmission. In humans, endogenous RNAi activity was reported in both germline and somatic cells^84–86^, although the interferon pathway is considered the main responder to dsRNA. Substantial evidence indicates that small RNAs originated from the mother can infiltrate the embryo and overtime effect global gene expression^71,87–89^. We hypothesize that the affected progeny may be continuously exposed to additional repeat-derived siRNAs during pregnancy. This may also explain the obscure disease course. Once the newborn overcomes the intensified RNA toxicity, the symptoms improve until the accumulative damage caused by the additional established RNA toxicity mechanisms is sufficient to produce classical DM1 symptoms. Importantly, the Maternal 123CUG nematodes exhibit evident maternal toxicity without anticipation of repeat length. Our suggested mechanism of maternally inherited toxicity suggests that repeat-carrying mothers may affect both sick and healthy progeny. Intriguingly, even healthy offspring born to DM1 mothers following pre-implantation genetic diagnosis presented a relatively high incidence of minor anomalies^90^. While this may be a result of existing medical problems in the DM1 mothers, our data suggest that siRNAs may play a role. The repeat siRNAs from the mother trigger abnormal gene silencing in these offspring, but the effect is moderate due to the lack of self-repeats. While maternal biases were characterized in DM1 and spinocerebellar ataxia type 8^7^, future research should investigate additional non-coding repeat expansion disorders. Moreover, such studies should evaluate the role of RNA interference in other disorders exhibiting parent-of-origin effects.

To conclude, we offer a novel mechanism of toxicity inheritance in repeat-based disorders. Our experimental data suggest that RNAi machinery may play a key role in the parent-of-origin effect and can transmit toxicity from parent to child, mediate disease pathogenesis, and explain the maternal bias. Consequently, it provides an opportunity to develop novel disease-modifying therapeutics for DM1 and possibly other expansion repeat disorders.

## Methods

### *C. elegans* and RNAi strains

*C. elegans* strains GR2024, GR3207 (123CUG) and GR2025, GR3208 (0CUG) were used^52^. These animals express 123CUG or 0CUG repeats in the 3’UTR of GFP in the body wall muscle cells under the myo-3 promoter. The N2 (Bristol) strain was obtained from the Caenorhabditis Genetics Center (Minneapolis, USA) and used as a wild-type strain. For the RNAi mutant assays, strains *ne299 (rde-4)*, and *PD8753 (dcr-1)* were obtained from the Caenorhabditis Genetics Center. *C. elegans* strains were handled using standard methods and grown at 20°C unless otherwise indicated^91^.

### Crossing Maternal and Paternal 123CUG strains

Ten 123CUG one-day-old males were put on a plate with four L4 wildtype hermaphrodites. Plates were synced after 48 hours, eggs were left to hatch overnight in M9, and L1 nematodes with Paternal 123CUG were produced. For Maternal 123CUG, the process was replicated with 10 wildtype one-day-old males and four L4 123CUG hermaphrodites. To rule out the possibility of biased phenotypes due to self-mating of the hermaphrodites, the crosses were validated by two approaches: ∼50% prevalence of males in the F1 offspring was assessed and randomly selected F1 animals were sequenced for presence of repeats.

### Gene inactivation

RNAi-mediated gene inactivation was achieved by feeding nematodes bacterial strains expressing dsRNA as previously described^92^. RNAi clones were obtained from the Ahringer’s library. A single colony of RNAi bacteria was grown overnight at 37°C in LB with 100 mg/ul ampicillin, and then seeded onto NGM plates with carbenicillin. Vector expression was induced by adding isopropyl β-D-thiogalactopyranoside (IPTG) for a final concentration of 0.5-1 mM directly over the bacterial lawn and left to dry for 24 h. An empty L4440 vector (EV) was used as a negative control.

### Motility

At day two, four, and six of adulthood, five *C*. elegans males were picked and placed on 60 mm NGM plates without food. The nematodes were left to recover for 20 minutes after which they were filmed. Over 15 animals were counted per experiment and the data from three biological replicates were combined. Images were captured using a digital microscope and Micam 4.0 Software. The resolution was 2048 × 1536 pixels and a total number of 120 frames were taken at a rate of 1 capture per second for 120 seconds. In each experiment, all images were captured with the same focus, on the same day, and at room temperature. The video was built by MakeAVi software with a playback rate of 15 frames per second. The animals were analyzed using Tracker 5.0 software by defining the tail of the animal as a point mass and manually tracking its position for each frame. The statistical analysis was performed using a one-tailed Student’s t-test, α = 0.05.

### Stress assays

Synchronized worm eggs were placed on NGM plates seeded with RNAi bacteria or OP50 (as indicated) and supplemented with 100 mM IPTG (4 mM final concentration). For the heat shock assay, at day one of adulthood 80 animals were transferred onto pre-warmed plates without bacteria (10 animals per plate) and exposed to 34-35°C. Survival rates were recorded every two h. Statistical analyses was performed using log-rank (Mantel-cox) and Gehan-Breslow-Wilcoxon tests, α=0.05.

### Morphology and length

Maternal and Paternal 123CUG animals were generated and left to hatch in M9 until the 1^st^ larval stage. Then they were placed on NGM plates seeded with OP50 and immediately imaged using a Nikon SMZ800N Microscope. After 72 h, the nematodes were imaged once again. The nematodes’ length was determined using the DeltaPix software. Statistical analyses were performed using a one-tailed Student’s t-test, α = 0.05.

### Target genes for siRNA silencing

A BLAST^93^ search was conducted to identify genes with seven or more CTG/CAG repeats and with no more than two mismatches, that could serve as the most obvious targets for siRNA silencing. Thirty-one transcripts were identified and specific primers for 24 of those genes were generated. Expression levels were determined using RT-qPCR as described below. We were unable to generate specific primers and establish expression levels for the seven remaining genes.

### RT-qPCR analysis of mRNA and siRNA expression

Total RNA was extracted from the whole body of *C. elegans* nematodes using Trizol Reagent (Ambion, USA) and a NucleoSpin RNA isolation kit (Macherey-Nagel, Germany). Hundreds of worms were collected for each experiment. For mRNA, reverse transcription was performed using a cDNA reverse transcription kit (Applied Biosystems, USA), and mRNA expression levels were measured with qPCR. SYBR-Green (Bio-Rad, USA) was used in a CFX-384 Real-Time PCR system (Bio-Rad). For siRNA, reverse transcription was performed using the MystiCq miRNA cDNA synthesis Mix (Sigma Aldrich, USA). MystiCq SYBR green (Sigma-Aldrich) and universal PCR primer (Sigma-Aldrich) were used for RT-qPCR. Data were analyzed using the ΔΔCt method. Relative quantities of gene transcripts were normalized to *rpl-32* and *cdc-42* for the mRNAs, and *mir-46-3p* for the siRNAs. All the primers used in this research were designed using the NCBI Primer Blast (sequences depicted in Table 2S).

### Fluorescence

Maternal 123CUG, Paternal 123CUG, and 0CUG nematodes were grown on empty vector bacteria and images were taken at the 1^st^ larval stage and two-day-old adults. The RNAi groups were treated as previously described and imaged at the same ages. The animals were washed twice with M9, anesthetized using 10 mM sodium azide (Sigma Aldrich, USA), and placed on an agar pad. Images were taken using a Spinning-Disk confocal microscope. For all fluorescence images, the images shown within the same figure panel were collected using the same exposure time and then processed identically in ImageJ. Statistical analyses were performed using a one-tailed Student’s t-test, α = 0.05.

## Data availability

The datasets generated during and/or analyzed during the current study are available from the corresponding authors on reasonable request.

## Acknowledgements

We thank Prof. Gary Ruvkun, Massachusetts General Hospital and Harvard Medical School, for his comments on the project. We also thank Prof. Igor Ulitsky and Prof. Eran Hornstein at the Weizmann Institute of Science for reviewing the manuscript. Funding was received from the Israel Science Foundation (grant agreement 1591/19).

## Author contributions

M.B. and Y.T. conceptualized the study and wrote the manuscript. M.B., S.S., J.T., A.M.S., and M.M. designed and carried out the experiments and analyzed the data. Z.G., S.E.J.F, S.M.D.A.G, and Y.T. edited the manuscript and supervised the research.

**The authors declare no competing interests**.

## Abbreviations

DM1: myotonic dystrophy type 1
DMPK: dystrophia myotonica-protein kinase gene
EV: empty vector (control RNAi)
OP50: *E. coli* strain
RNAi: RNA interference
3’UTR: 3’ untranslated region
wt: wild type

