## Supplemental files for "RNA interference mediates RNA toxicity with parent-of-origin effects in *C. elegans* expressing CTG repeats"

**Figure 1S:** Maternal origin of repeats reduces worm size

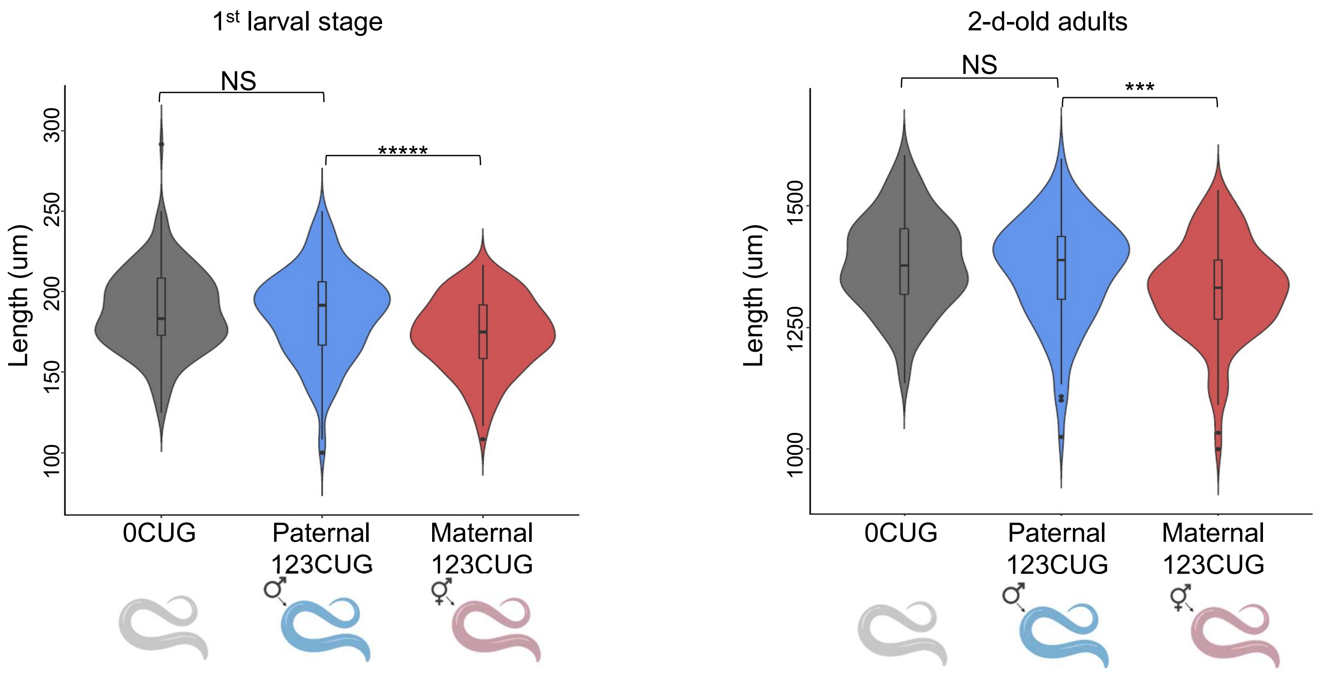

**Figure 2S:** Motility phenotype recapitulated on a different 123CUG strain

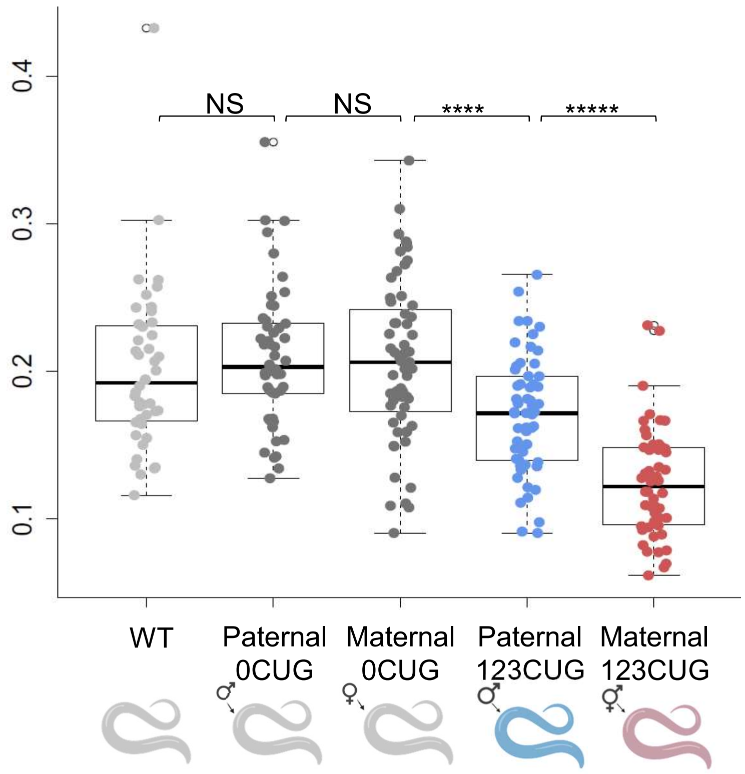

**Figure 3S:** Enrichment of CUG-repeat siRNAs in 123CUG groups

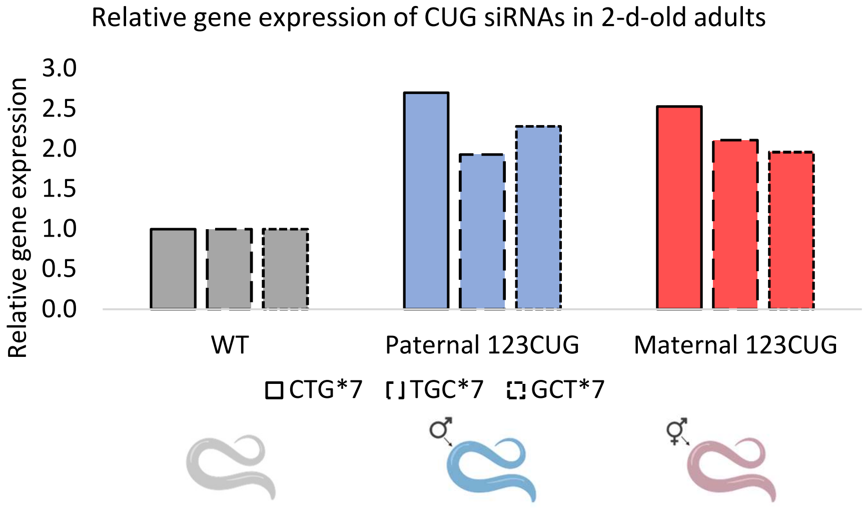

**Figure 4S:** *dcr-1* and *rde-4* knockout mutants present with severely impaired motility

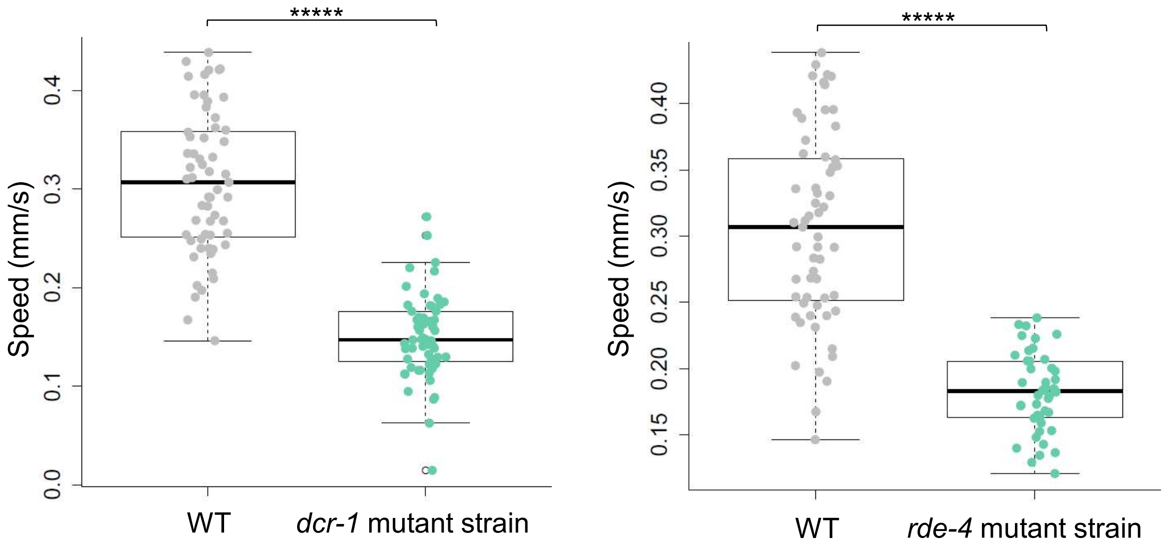

**Figure 5S:** (A) *dcr-1* RNAi does not affect 0CUG and Paternal 123CUG motility

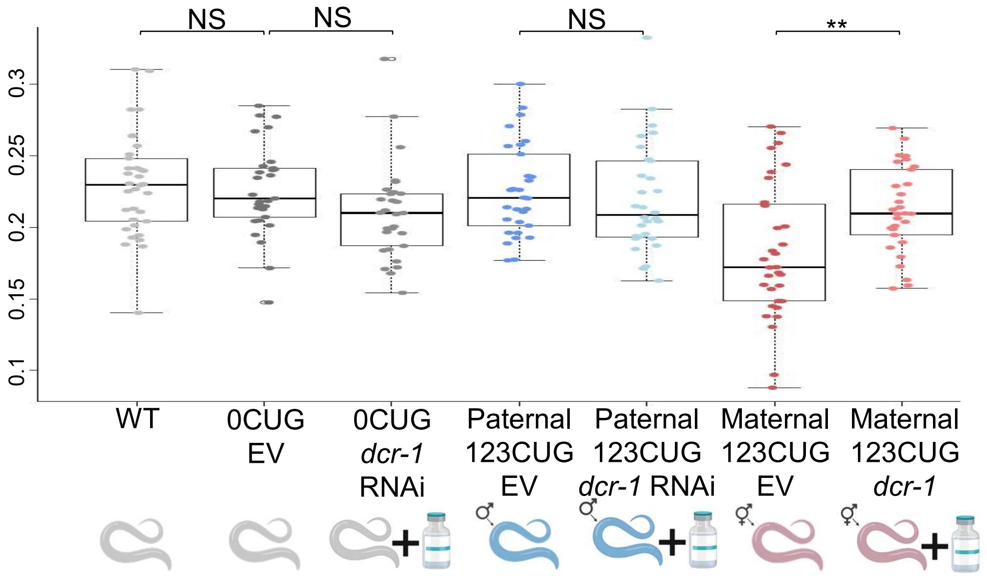


(B) *rde-4* RNAi does not affect 0CUG and Paternal 123CUG motility

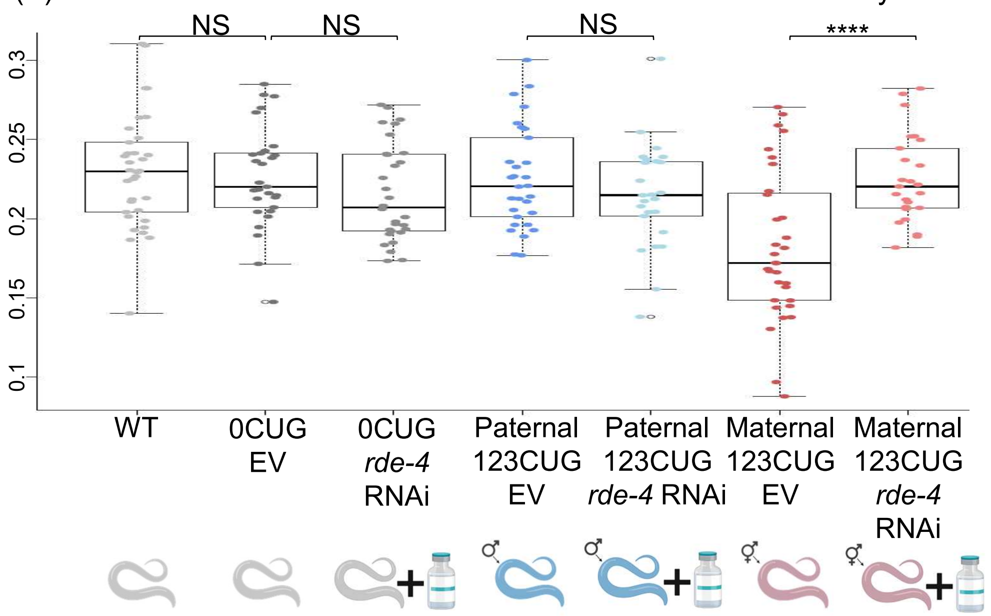


(C) *rde-1* RNAi does not affect 0CUG and Paternal 123CUG motility

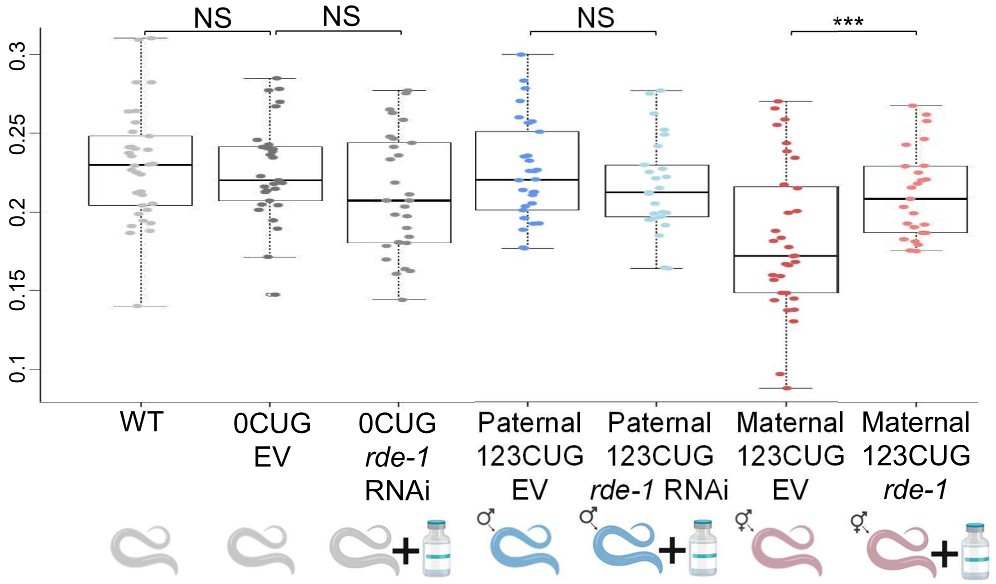

**Figure 6S:** Fluorescence of Paternal 123CUG and treatment groups did not differ

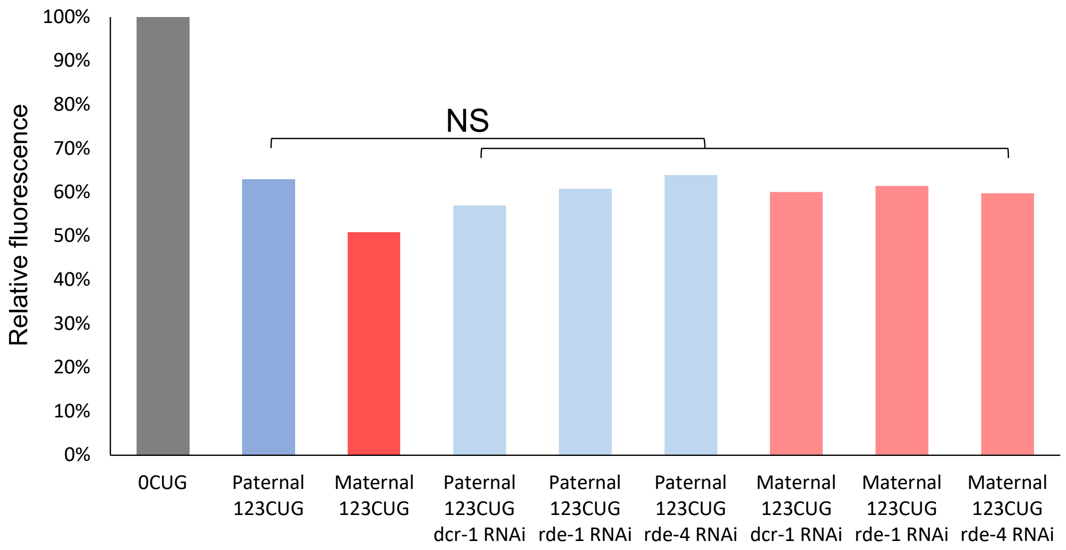

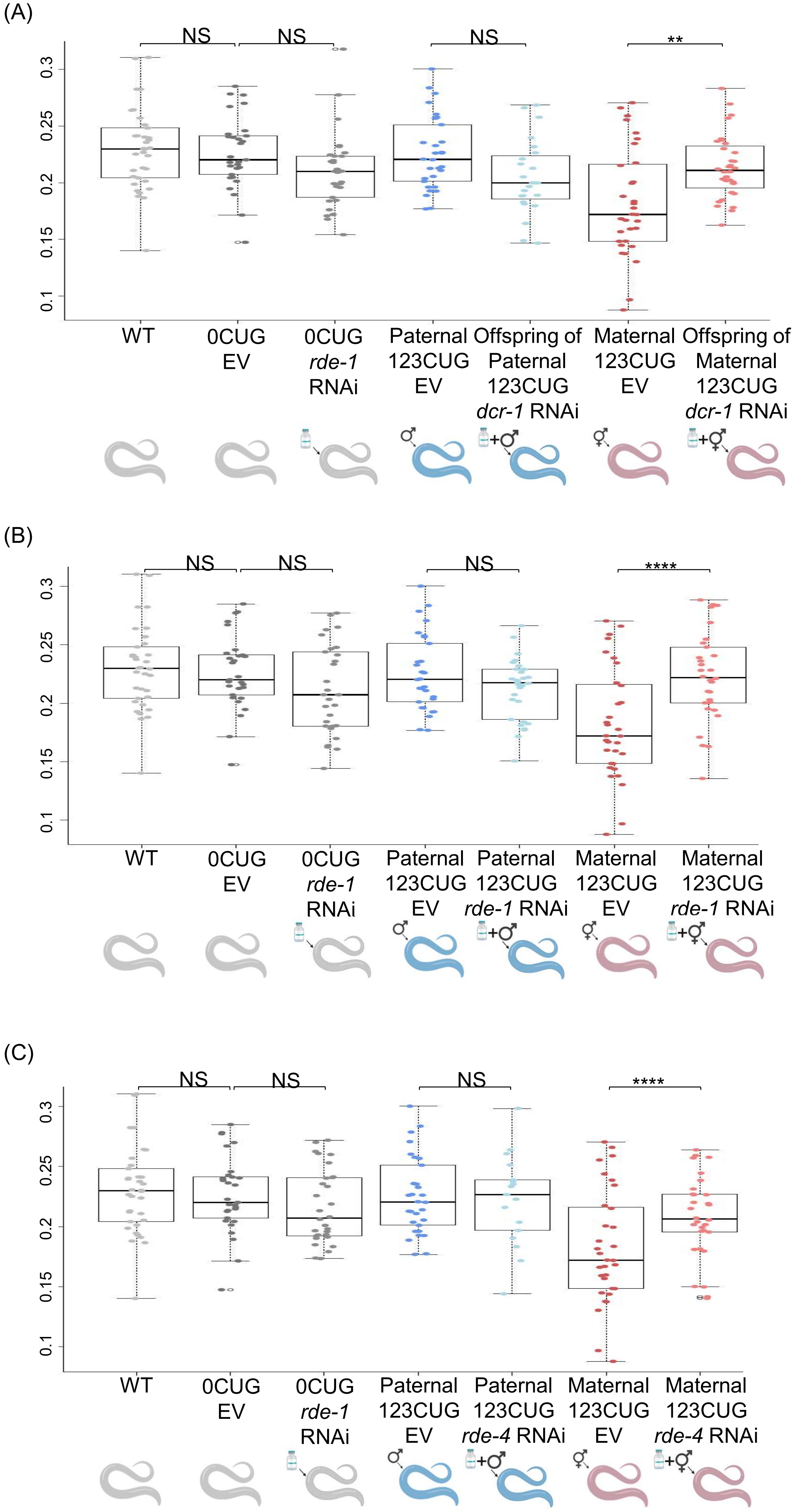

(A)
NS
NS
NS
**
0.3
0.25
0.2
0.15
0.1
WT
0CUG EV
0CUG *rde-1* RNAi
Paternal 123CUG EV
Offspring of Paternal 123CUG *dcr-1* RNAi
Maternal 123CUG EV
Offspring of Maternal 123CUG *dcr-1* RNAi
(B)
NS
NS
NS
****
0.3
0.25
0.2
0.15
0.1
WT
0CUG EV
0CUG *rde-1* RNAi
Paternal 123CUG EV
Paternal 123CUG *rde-1* RNAi
Maternal 123CUG EV
Maternal 123CUG *rde-1* RNAi
(C)
NS
NS
NS
****
0.3
0.25
0.2
0.15
0.1
WT
0CUG EV
0CUG *rde-1* RNAi
Paternal 123CUG EV
Paternal 123CUG *rde-4* RNAi
Maternal 123CUG EV
Maternal 123CUG *rde-4* RNAi

**Figure 8S:** No difference in fluorescence observed between Paternal 123CUG group and treatment groups

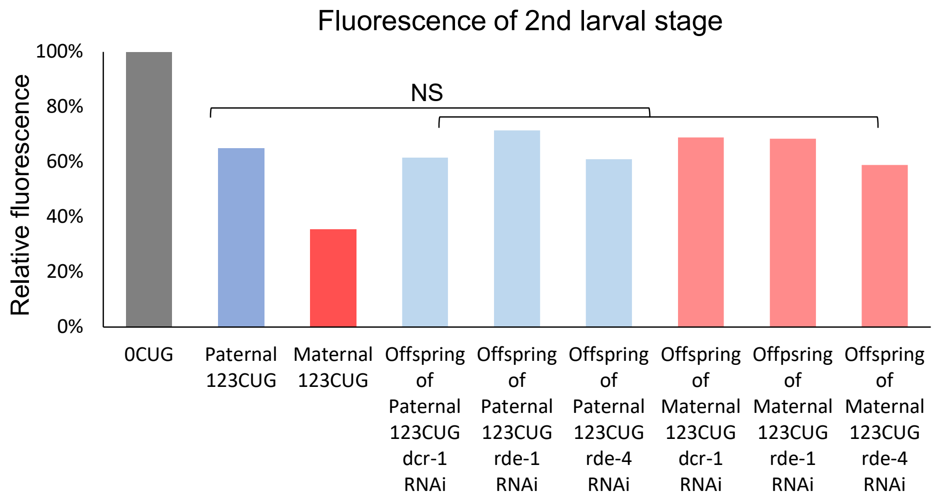


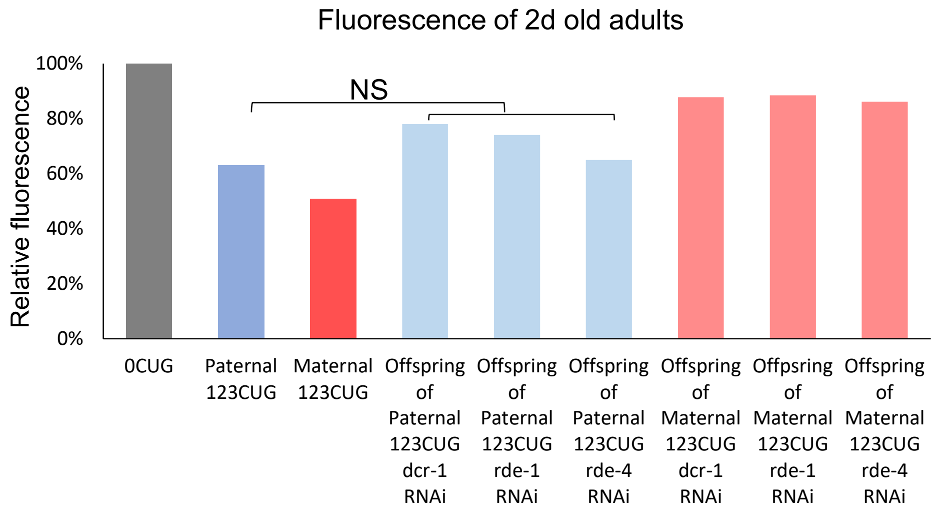
